# Nanobody MET CAR-T cells show efficacy in solid tumors

**DOI:** 10.64898/2026.01.27.702111

**Authors:** Po-Han Chen, Qin Li, Sam Deveraux, Danielle K. Sohai, Pei-Chun Cha, Rianna Raghunandan, Nancy Chen, Yao Li, Mileena Nguyen, Michael C. Stankewich, Jon S. Morrow, Arnaud Augert, Qin Yan, Samuel G. Katz

## Abstract

**Background:** MET overexpression is associated with poor prognosis in many solid tumors due to its central role in tumor survival, invasion, metastasis, and chemoresistance. While targeting MET with antibody-drug conjugates has shown promising results, engineered cellular immunotherapeutic approaches have not been extensively explored. Compared to conventional single-chain variable fragments (scFv), naturally occurring single-domain antibodies consisting of variable heavy chains only (VHH or nanobodies) are smaller, retain high specificity, and exhibit remarkable biochemical stability. In this study, we tested the efficacy of MET-targeting VHH-CAR-T (chimeric antigen receptor T cells).

**Methods:** We generated a panel of VHH-CAR-Ts using mRNA electroporation. VHH-CAR-T cells were evaluated in functional assays including cell binding avidity, cytokine production profiles, hydrogel microwell-based cellular kinetics, and in vitro cytotoxicity. We also assessed the therapeutic efficacy of VHH-CAR-T in an in vivo mouse model of metastatic triple negative breast cancer (TNBC).

**Results:** Among the tested VHH, we identified those with intermediate avidity as most effective for in vitro tumor killing. VHH-CAR-Ts with CD28 costimulatory domains demonstrated augmented cytotoxicity with favorable selectivity, requiring a minimum antigen density threshold for activation. Mechanistically, VHH-CAR-Ts demonstrated low tonic signaling, high avidity, potent cytokine production, and rapid tumor killing kinetics. When administered in an mRNA format, VHH-CAR-Ts exhibited potent and prolonged control of tumor growth in an in vivo metastatic model of TNBC.

**Conclusion:** Taken together, these results demonstrate that VHH-CAR-Ts exhibit robust MET specificity and potent therapeutic efficacy both in vitro and in vivo. Thus, VHH-CAR-T cell therapy represents a promising immunotherapeutic strategy for targeting MET-overexpressing solid tumors.

**What is already known on this topic:** MET signaling is an important contributor to the aggressiveness of many solid tumors, and targeting MET by antibody-drug conjugates has shown efficacy and safety. Targeting MET by CAR-T cells has been under study, though with limited potency.

**What this study adds:** This study is the first to demonstrate effectiveness of anti-MET VHH-CAR-T cells. Compared with other antigen binding domains, VHH-incorporated CAR-T cells show low tonic signaling, a favorable cytokine profile, and potent tumor killing.

**How this study might affect research, practice or policy:** With the multiple advantages of VHHs including small size, stability, and low potential for tonic signaling, VHH-CAR-T cells represent a promising approach for CAR-T design against solid tumors.

## INTRODUCTION

While chimeric antigen receptor (CAR) engineered T cells have shown impressive efficacy in hematologic malignancies, solid tumors pose many challenges. Target selection remains a challenge for solid tumors due to the lack of tumor-specific antigens[1]. Innovative CAR designs are also needed to promote tumor infiltration and persistence in the hostile tumor microenvironment, to overcome heterogeneous antigen expression, and to avoid on-target/off-tumor cytotoxicity.

The *MET* oncogene encodes the MET receptor tyrosine kinase, which is overexpressed in many solid tumors and is associated with poor prognosis due to its central role in mediating drug resistance and pro-survival signaling[2]. While tyrosine kinase inhibitors are available, their efficacy is limited to a select number of MET-addicted tumors with MET amplification or mutations[2]. In contrast, targeting MET expression with surface antibodies such as antibody-drug conjugates (ADCs) has shown safety and copy number/mutation-independent efficacy[3]. As CARs leverage the antigen recognition ability of antibodies with the additional benefits of adaptive T cell responses, targeting MET overexpression using CAR-T cells holds great potential.

The antigen recognition domain affects the target sensitivity, durability, and tonic signaling of CAR-T cells[4,5]. Most CARs use a single-chain variable fragment (scFv) as the antigen binding domain, which commonly exhibits self-clustering and tonic signaling leading to T cell exhaustion[6,7].

A variable region of a heavy-chain-only antibody (VHH or nanobody) is emerging as an alternative to scFv[8–10]. VHH-based CARs have been successfully used in BCMA-CAR-T cell therapy ciltacabtagene autoleucel (cilta-cel) against myeloma with approval from the U.S. Food and Drug Administration (FDA) [11]. Of particular importance for solid tumors, the stability of VHH helps avoid receptor clustering and tonic signaling; its uniquely long complementarity-determining region 3 (CDR3) enables targeting of cryptic epitopes and isoforms that may overcome tumor heterogeneity; its compact size allows flexible integration with other moieties (such as tandem VHH in cilta-cel) to facilitate innovative CAR designs with enhanced functionality; finally, its high degree and ease of humanization reduce immunogenicity and ensure safety in humans[8,11–15].

In this study, we constructed a panel of mRNA VHH-CAR-T cells targeting MET, and demonstrate that VHH-CAR-T cells exhibit high avidity, potent cytokine production upon stimulation, a potential therapeutic window for CAR activation, low tonic signaling, and effective control of metastatic tumors in vivo – features that warrant further exploration as a novel form of T cell immunotherapy for solid cancers.

## MATERIALS AND METHODS

### Cells cultures

Breast cancer cell line MDA-MB-231 was purchased from ATCC (HTB-26), and its metastatic sub-population LM2 was a kind gift from J. Massagué. The non-small cell lung cancer cell line A549 cell line was obtained from Peter Glazer, and the head and neck squamous cell carcinoma cell lines FaDu and CAL27 cell lines were obtained from Barbara Burtness. These cells were cultured in DMEM+10% FBS+1% penicillin/streptomycin. The renal cancer cell line 786-O was obtained from Harriet Kluger and cultured in RPMI1640+10% FBS+1% penicillin/streptomycin. The diffuse large B-cell lymphoma cell lines Karpas-422 and OCI-LY3 were obtained from Markus Müschen and cultured in RPMI1640+20% FBS+1% penicillin/streptomycin. Small-cell lung cancer cell lines NCI-H146 (ATCC HB173) and NCI-H1341(ATCC CRL-5864) were cultured in DMEM-F12 supplemented with 10% FBS, 1X Insulin-Transferrin-Selenium and 1% penicillin/streptomycin.

### Construction of CAR and *in vitro* transcription

The CAR construct consists of a T7 promoter, 5’UTR (human beta-globin), CAR, P2A self-cleavable sequence, GFP, and 3’UTR (human beta-globin) (see **Supplemental Figure 1A**). Binding domains for the nanobodies (VHH1-5), 1E4 (scFv1) and 5D5 (scFv2) sequences were adapted from patents WO2012/042026[16], US10116622B2[17] and US20190085063A1[18], respectively. The scFv3 and NK1-CAR sequences were adapted from previous publications [19–21]. The CAR sequences were synthesized as gene blocks containing T7 promoter and 5’ and 3’ UTR sequences. They were cloned into standard pUC57 plasmid using the InFusion cloning kit (Takara).

For *in vitro* transcription, a PCR product encompassing the T7 promoter and 3’ UTR, where the reverse primer also added a 100 nucleotide poly-A tail[22], was used in the T7 ARCA HighScribe mRNA Kit with Tailing Reaction (NEB, Cat. #E6020S). Final mRNA was purified using lithium chloride precipitation and concentrated to 1-2 μg/μL in water.

### Isolation and activation of primary human T cells

Human peripheral blood mononuclear cells were collected from healthy donors (either local draws under IRB protocol HIC2000033275 or from the Gulf Coast Regional Blood Center). Bulk T cells were isolated using the EasySep Human T Cell Enrichment Kit (STEMCELL Technologies, Cat. #17951), activated with ImmunoCult Human CD3/CD2/CD28 T cell activator (STEMCELL, Cat. #10990) at the recommended concentration, and cultured at 1 x 10^6^ cells per mL in RPMI1640 + 10% FBS in the presence of 100 IU/mL recombinant human IL-2 (Miltenyi Biotec, Cat. #130-097743).

### Production of mRNA CAR-T cells

On days 7-10 post T cell activation, T cells were pelleted, washed, and resuspended in 20 μL MaxCyte buffer per 2 million cells. Electroporation was performed using the MaxCyte ATX with the Expanded T cell Protocol 1 at 1 μg mRNA/1 million T cells. Following electroporation, cells were rested in a 37^0^C incubator with 5% CO_2_ for 10 minutes and resuspended in complete T cell culture media at 2×10^6^ cells/mL with 100 IU/mL IL-2. Cells were either frozen after 4 hours or used for experiments after overnight incubation. Successful mRNA electroporation was confirmed by GFP expression using either fluorescence microscopy or flow cytometry.

### Flow cytometry

Immunophenotypes of all samples were analyzed using CytoFLEX S (Beckman Coulter, CA, USA) flow cytometers. FlowJo software (Tree Star, OR, USA) was used for data analysis. The following antibody-conjugated fluorophores were used: CD3 AlexaFlour 700 (1:100, clone OKT3, BioLegend 317340); CD8 BUV711 (1:100, clone RPA-T8, BioLegend 301044); CD4 BUV605 (1:100, clone OKT4, BioLegend 317438); CD69 PE/Cy5 (1:100, clone FN50, BioLegend 310908); TNF-α PE (1:50, clone MAb11,BioLegend 502909); IFN-γ PerCP/Cy5.5 (1:100, clone XMG1.2, BioLegend 505822); IL-2 BV785 (1:50, clone MQ1-17H12, BioLegend 500348); IL-4 PE/Dazzle 594 (1:100, clone MP4-25D2, BioLegend 500832); PD-1 PE/Cy7 (clone A17188B, BioLegend 621616); TIM-3 APC (clone A18087E, BioLegend 364804); LAG3 APC/Fire750 (clone 7H2C65, BioLegend 369214). MET expression was confirmed by flow cytometry with a monoclonal anti-MET antibody (1:10, Roche, clone SP44) compared to an IgG isotype control (BioLegend 400120).

### Gene expression database analysis

Gene expression data for MET across human small cell lung cancer (SCLC) cell lines were obtained from the SCLC-CellMiner database using the SCLC Global response cell line set[23]. Z-scores corresponding to MET expression values were extracted and visualized as a heatmap using GraphPad Prism to compare relative expression levels across cell lines.

### Western blot

Protein samples were resolved on 4–20% Mini-PROTEAN TGX precast polyacrylamide gels (15-well, 15 µl; Bio-Rad, #4561096) and transferred to nitrocellulose membranes. Membranes were blocked in 5% non-fat milk in Tris-Buffered Saline with 0.1% Tween-20 (TBST) and incubated with primary antibodies overnight at 4°C (Beta-Actin, 1:5,000, GTX629630; MET, 1:1,000, Cell Signaling Technology #4560). After washing, membranes were incubated with HRP-conjugated secondary antibodies (1:3,000; Cell Signaling Technology; anti-rabbit IgG #7074, anti-mouse IgG #7076) in 5% non-fat milk in TBST for 2 hours at room temperature. Blots were developed using an ultra-sensitive enhanced chemiluminescent (Thermo Scientific, #34095) and imaged on Bio-Rad ChemiDoc imaging system.

### Immunohistochemistry (IHC)

4-µm sections were obtained from formalin-fixed, paraffin-embedded (FFPE) tissue. Anti-MET antibody (clone SP44, AbCam, ab227637) was manually applied at 1:50 dilution after antigen retrieval using Tris-EDTA buffer at pH 9.0. Anti-human CD3 antibody (Biocare Medical, CP215) was applied at 1:75 dilution manually following vendor protocol. IHC quantification was performed by visual examination of multiple high power fields for % tumor positive for MET (all intensity).

### CD69 activation assay for CAR-Jurkat

The protocol was similar to that previously described [24]. Briefly, 30,000 target cells per well were plated in a 96-well plate. Jurkat cells were electroporated with CAR mRNA and incubated at 37^0^C for 4 hours before adding to target cells at various E:T ratios (no target, 100:1, 10:1, 1:1). After overnight incubation, Jurkat cell activation was measured by surface expression of CD69 using flow cytometry.

### In vitro tumor killing assays

The flow-based killing assay was similar to that previously described[25] with the following modification. Tumor cells labeled with 1 μM CellTrace Far Red (Invitrogen C34572) were seeded in 24-well plates at a concentration of 1 x 10^5^ cells/well. Mock or CAR-T cells were added to the culture at different ratios (E:T of 0.5:1, 1:1, 2:1, 4:1) without adding exogenous cytokines. Cells were analyzed 16 hours later to measure residual tumor cells by flow cytometry. Dead cells were identified by Live/Dead Violet staining (Invitrogen L34955). Tumor killing was calculated using the formula: % specific lysis = (1-# residual target cells in treated group/# target cells cultured alone without T cells) x100%.

The luciferase-based killing assay was performed using cells expressing firefly luciferase (MDA-MB-231, FaDu, and CAL27). Tumor cells were plated with CAR-T cells at the desired E:T ratio in 384-well plates in 30 μL media overnight. Luminescence was detected using the ONE-Glo Luciferase Assay System (Promega E6120) and analyzed using a plate reader (BioTek, SYNERGYMx). Cell lysis (%) = (1 – (experimental readout – spontaneous readout)/spontaneous readout) x 100.

### CellCyte tumor proliferation assay

A lentiviral plasmid encoding mKate2 with puromycin selection was obtained from Dr. David Fox at the University of Michigan. A549 tumor cells expressing mKate2 were co-cultured with T cells in 96-well optically clear flat-bottomed plates at an E:T ratio of 3:1. Plates were imaged using the CellCyte system (ECHO) every 4 hours, and the number of mkate2-positive tumor cells was counted by the provided software.

### Mouse tumor xenograft

Six-week-old female NOD.Cg-Prkdc^scid^ Il2rg^tm1Wjl^/SzJ (NSG) mice were purchased from the Jackson Laboratory (Strain #005557). Tumor inoculation was performed via tail vein injection of 2×10^5^ cells in 100 μL PBS when the mice reached 9-10 weeks of age. Once injected tumors reached a total flux between 10^7^ to 10^8^ photons/sec by bioluminescence imaging (BLI) (∼7-10 days), mice were randomized into respective groups and given weekly injections of PBS or T cells as specified in the text. To avoid graft-versus-host disease, the day before each subsequent T cell injection, 60 mg/kg cyclophosphamide was injected intraperitoneally to deplete the previously injected T cells [26]. Tumor burden was monitored by weekly BLI using the Xenogen *In Vivo* Imaging System (IVIS) (Caliper Life Sciences) and analyzed with Living Image software. Animals were euthanized once luminescence signals reached 1×10^9^ photons/sec or when animals developed significant stress.

### Cell force binding assay (avidity)

CAR-tumor binding force was measured using the z-Movi device (LUMICKS, Amsterdam, The Netherlands). Target cells were seeded onto microfluidic chips at a concentration of 5×10^7^ cells per mL to obtain a confluent monolayer. CAR-T cells were labeled with 1 μM CellTrace Far Red to enable T cell tracking. After co-incubation of CAR-T cells with the tumor monolayer for 2.5-5 minutes, a force ramp ranging from 0 to 1000 pN was applied for 2.5 minutes. T cell detachment was detected by fluorescence imaging. The binding force was analyzed using the Oceon software (V.1.0, LUMICKS).

### Cytokine release assay

For intracellular cytokine staining assays, T cells and target cells were co-cultured at a 1:1 E:T ratio for 5-12 hours in the presence of 2 μM brefeldin A (BD GolgiPlug, cat # 51-2301KZ). For the bulk cytokine assay, the supernatants from overnight co-culture were harvested and stored in –20^0^C before shipping to Eve Technologies.

For the multiplexed secretome assay, after stimulation with target cells, approximately 30,000 magnetically enriched CD8+ CAR-T cells were processed for membrane staining (AF647-CD8, STAIN-1002-1). Subsequently, these cells were loaded onto the IsoCode chip (IsoPlexis, ISOCODE-1001-4), comprising 12,000 chambers prepatterned with an array of 32 cytokine capture antibodies. The chip was then incubated in the IsoLight machine for 16 hours at 37°C with 5% CO_2_ supplementation. A cocktail of detection antibodies was then applied to detect the secreted cytokines, followed by fluorescence labeling. The resulting fluorescence signals were analyzed using IsoSpeak v.2.8.1.0 (IsoPlexis) to determine the number of specific cytokine-secreting cells and the intensity level of each cytokine.

### Structural modeling

The structures of the VHH sequences, scFv1 (1E4) and scFv2 (5D5), and their respective complexes with the human MET extracellular domain were predicted using AlphaFold3[27]. Antibody-MET affinity was estimated using the HADDOCK PRODIGY server [28,29]. Structural modeling and analysis were performed using PyMOL (Schrödinger, LLC).

### Microwell co-culture system

We used the TROVO system (Enrich Biosystems) for the microwell assays. For the tumor killing and T cell proliferation assay, T cells (stained with CD4/CD8-Alexa647) and MDA-MB-231 cells were plated at a 2:1 effector to target ratio in a six-well culture plate, with each well containing 3,200 printed hydrogel microwells. Each microwell was indexed by proprietary software and imaged every 12 hours over a four-day period for downstream analysis. Tumor killing was defined as a ≥15% reduction in tumor area compared to day 0. T cell proliferation was defined as a ≥2-fold increase in stained T cell area compared to day 0. For the single cell killing assay, CAR-T cells and tumor cells were plated at a 1:10 E:T ratio, and only microwells containing a single T cell were selected for further imaging analysis. The percentages of microwells with tumor killing/T cell proliferation, and rate of killing were calculated by the proprietary software built into the TROVO system.

In detail, percentage of microwells with tumor killing or T cell proliferation is calculated as number of microwells meeting the killing/proliferation threshold stated above divided by total number of microwells in this well. Bar graph is average +/− standard deviation of 3 technical replicate wells.

Single cell killing rate was estimated by fitting the paired T cell and tumor signals (area or counts) at each time point to a mass-action interaction model. For each CAR-T construct (and matched controls), we ran three independent 3,200-well microwell plates (9,600 wells/clone total). Dilute seeding targeted ∼1 T cell and ∼10 tumor cells per well. Plates were imaged from 0–96 h; fluorescence at time zero was used to calibrate area-to-cell conversions. Tumor-only wells on the same plates were fit to an exponential model C(t) = C₀e^[rC·t] to estimate the plate-level tumor growth rate rC. From the full set, we retained ∼800–900 co-culture wells per sample that met pre-specified QC (confirmed single T at time zero, adequate tumor baseline, stable focus/tracking, and finite parameter estimates). An effective killing rate (k_eff) was then obtained per well using a simplified Lotka–Volterra–type predator–prey coupling dC/dt = rC·C − k_eff·T·C (classical LV prey equation) and the same mass-action kill term described in similar CAR-T models[30], i.e., tumor loss regressed against cumulative T cell exposure. Fits were performed by nonlinear least squares with plate-specific rC; Fitting results with goodness-of-fit R² > 0.95 are included. For each CAR-T construct, the distribution of k_eff was summarized per plate, and the construct-level average k_eff and standard deviation was calculated across the triplicated experiments.

### Statistics

All statistical analyses were conducted using Prism v.10 (GraphPad). Unless otherwise specified, data were presented as mean ± SD. Statistical significance was determined by paired t-test, one-way analysis of variance (ANOVA), or two-way ANOVA as indicated in the figure legends. The log-rank (Mantel-Cox) test was used to determine statistical significance for overall survival in mouse experiments.

## RESULTS

### VHH-CAR-T with intermediate avidity showed the most potent tumor killing

We curated five camelid-derived VHH nanobody sequences and incorporated them into a second-generation mRNA CAR design with a CD8 hinge and transmembrane domain, followed by 4-1BB and CD3ζ signaling domains (**Supplemental Figure 1 and Figure 1A**).

**Figure 1.**
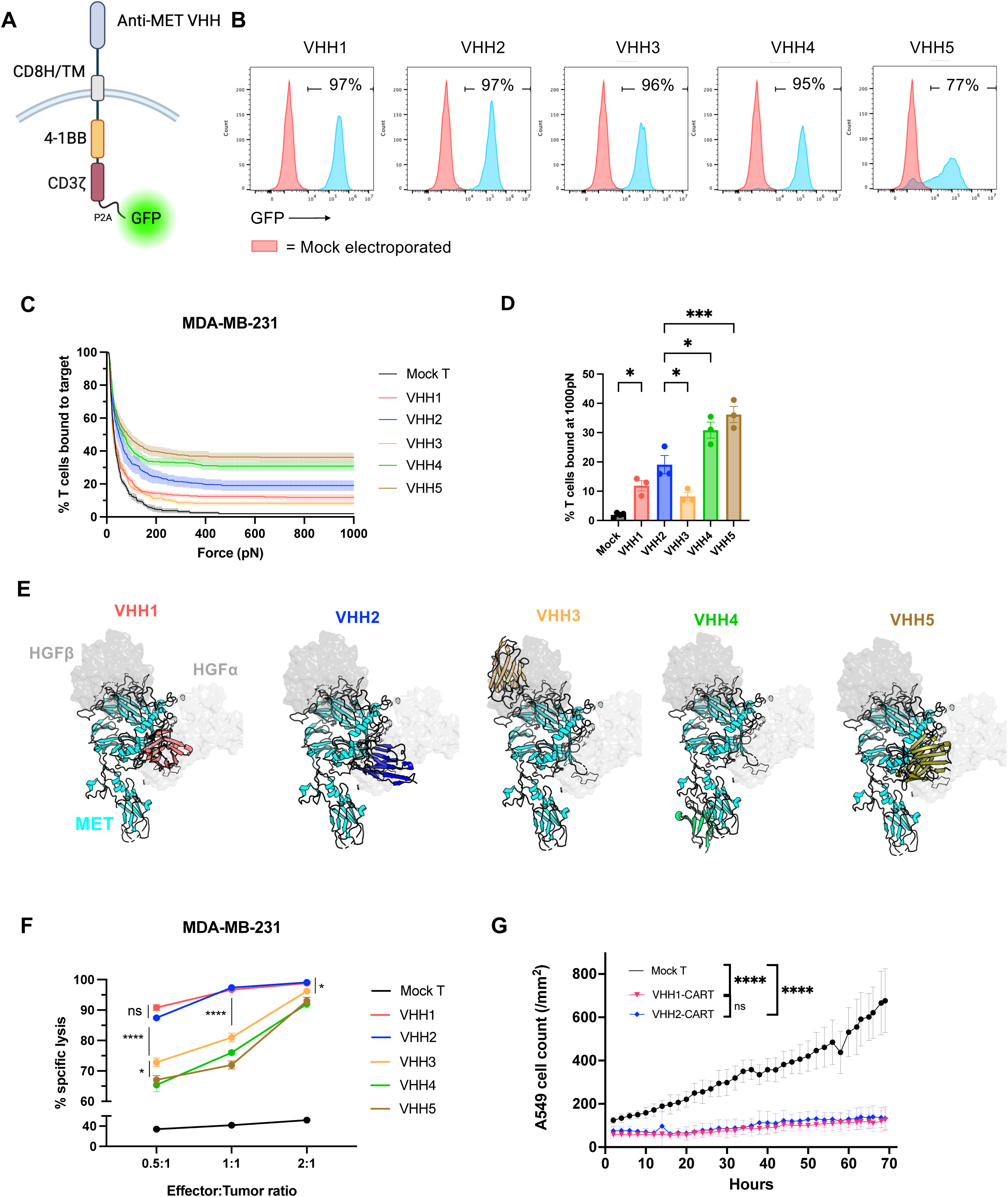
VHH-CAR-T with intermediate avidity showed the most potent killing. (A) Schematic diagram of VHH-CAR construct. (B) Expression level of VHH-CAR by GFP. Red peak = mock electroporated T cells. (C) Avidity curve of VHH-CAR-T against MDA-MB-231, with the % T cells bound to target cells at 1000 pN shown in (D). N = 4 technical replicates; *P* values comparing avidity at 1000 pN were calculated using one-way ANOVA). (E) AlphaFold3 modeling of VHH binding sites on MET. MET = cyan; HGFα chain = light gray; HGFβ chain = dark gray; VHH1 = red; VHH2 = blue; VHH3 = orange; VHH4 = green; VHH5 = brown. (F) VHH-CAR-T cytotoxicity against MDA-MB-231 (n=4 technical replicates; *P* values by two-way ANOVA). (G) VHH1 and VHH2-CAR-T against A549, with tumor proliferation monitored in CellCyte (n=6 technical replicates; *P* values by two-way ANOVA). Data shown are mean±SD; \**P<0.05; **P<0.01; ***P<0.001; ****P<0.0001*.

Since avidity represents the strength of the immune synapse and has been shown to predict clinical response[31], we used the z-Movi system to measure the avidity of different VHH-CAR-T cells. To ensure that avidity measurement was not confounded by different CAR expression levels, the amount of electroporated mRNA was adjusted so that all constructs had >85% expression as determined by GFP; however, VHH5-CAR was consistently expressed at ∼75-80% regardless of amount of mRNA input (**Figure 1B**).

All VHH-CAR-Ts showed varying percentages of bound T cells with increasing rupture force (**Figure 1C**). VHH1, VHH2, and VHH3 showed relatively low avidity compared to VHH4 and VHH5. Despite lower expression, VHH5 surprisingly showed the highest avidity. The mean percentage of remaining T cells bound to targets at 1000 pN, from the lowest to highest, was VHH3 (7%), VHH1 (12%), VHH2(19%), VHH4 (31%), and VHH5 (36%) (**Figure 1D**).

To gain structural insight into functional avidity, we modeled the binding of the different VHHs to MET using AlphaFold3 (**Figure 1E**). VHH1, VHH2, and VHH5 are predicted to overlap with the high affinity binding site for the HGF-α chain. VHH3 is predicted to bind to a more membrane-distal epitope that overlaps with HGF-β chain. VHH4 appears to bind to a distinct epitope separate from the other VHHs, slightly away from either HGF binding site. These predictions are consistent with the reported antagonistic effect of these VHHs on the HGF-MET interaction. The dissociation constant *K*_D_ was estimated by PRODIGY for the top-ranked structures. The affinity for VHH3 ranged from ∼10-50 nM; VHH1 and VHH2 showed intermediate affinity, averaging 5-10 nM; VHH4 and VHH5 had the highest affinity, ranging from 0.4-1.2 and 2-3 nM, respectively. The estimated affinities correlated with the observed avidity, suggesting a direct relationship between affinity and immune synapse strength.

We then investigated whether different avidities correlate with the in vitro cytotoxic functions of effector cells, expecting higher avidity to achieve better killing of MDA-MB-231 cells. Surprisingly, we found that lower avidity CARs mediated stronger tumor killing, with VHH1 and VHH2 achieving the highest level of cytotoxicity (**Figure 1F**). VHH3 showed the lowest avidity, but achieved slightly better killing than VHH4 and VHH5. The superior tumor killing by VHH1, VHH2, and VHH3 was most notable at low E:T ratios. There was no significant difference in cytotoxicity between VHH1 and VHH2. Consistently, we showed that both VHH1 and VHH2 strongly inhibit proliferation of A549 lung cancer cells over a 70-hr period using the CellCyte system (**Figure 1G**).

Although high-avidity CARs are generally favored and have been shown to correlate with clinical response, our findings are consistent with emerging evidence documenting potential advantages of lower-avidity CARs (see Discussion). Given that there was no significant difference between VHH1 and VHH2 in terms of avidity and in vitro cytotoxicity, we used VHH2 for further characterization.

### CD28 costimulatory domain conferred stronger VHH-CAR-T activation than 4-1BB

Using VHH2, we next investigated how different cytoplasmic costimulatory domains, namely, CD28 versus 4-1BB, affect VHH-CAR-T function. Although not significantly different, VHH2-28z trends towards higher avidity than VHH2-BBz (**Figures 2A and B**). VHH2-28z also achieved higher in vitro tumor killing and more immune cytokine (IFN-γ, TNF-α, IL-2, and IL-4) and cytotoxic molecule Granzyme B production, as well as higher polyfunctionality (**Figures 2C-E, and Supplemental Figure 2**). In addition to these fixed-time point assays, we also used a microwell-based co-culture platform to gain insights into the kinetics of CAR-T cell killing. In microwells where we seeded 1 effector T cell per 10 tumor cells (1:10 E:T), we found that VHH2-28z demonstrated a higher single-cell killing rate than VHH2-BBz (**Figure 2F**). At an experimental setup of a 20:10 E:T ratio, we observed more tumor killing and higher T cell proliferation associated with VHH2-28z compared to VHH2-BBz over a four-day period (**Figures 2G-I**). Although CD28 costimulatory domain confers immediate potency, it is reported to be more prone to exhaustion than 4-1BB[32]. However, most likely due to the transient nature of mRNA CAR expression, we did not observe significant increase in exhaustion markers PD-1, LAG3, or PD-1+/TIM3+ for VHH2-28z (**Supplemental Figure 3**).Our data is consistent with higher and more rapid cytotoxic function generally reported for CD28 compared to 4-1BB[32,33]. Beyond four days, most likely due to diminishing mRNA, there was no significant killing or proliferation observed with either construct (data not shown). In subsequent studies, we used VHH2-28z unless otherwise specified.

**Figure 2:**
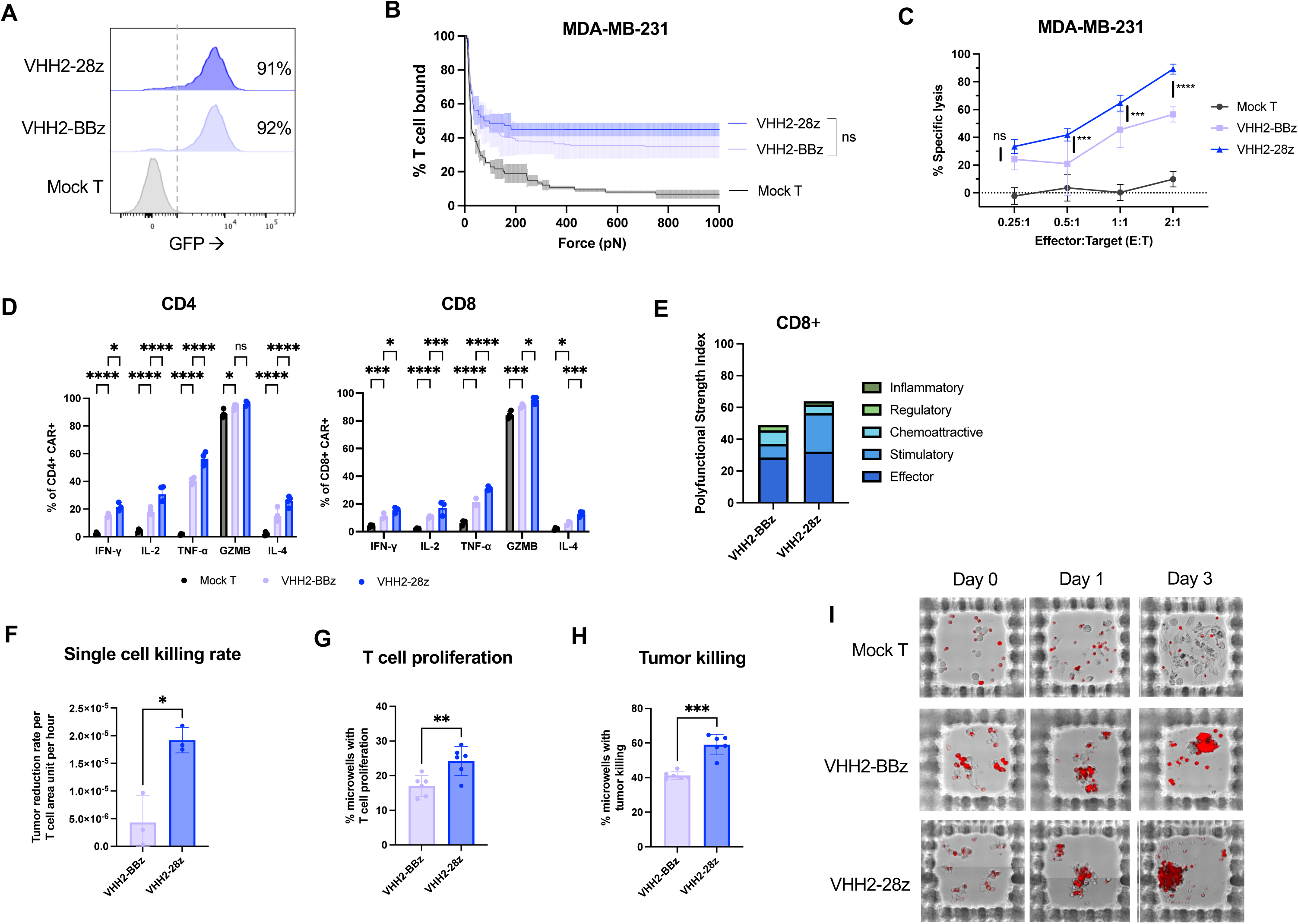
CD28 costimulatory domain conferred stronger VHH2-CAR activation. (A) Expression level of mRNA VHH2-BBz and VHH2-28z in primary human T cells. (B) Avidity assay using Lumicks z-Movi, comparison between VHH2-BBz and VHH2-28z (n = 3 technical replicates; two-way ANOVA). (C) Comparison of MDA-MB-231 killing between VHH2-BBz and VHH2-28z (n=8 technical replicates, two-way ANOVA). (D) Intracellular cytokine production analyzed by flow cytometry; cells harvested 16 hours after co-culture with MDA-MB-231 in the presence of 2μM of the Golgi inhibitor monensin (n = 4 technical replicates, two-way ANOVA). (E) Polyfunctioinal strength index (PSI) using IsoPlexis, comparison between CD8+ VHH2-BBz and CD8+ VHH2-28z 16 hours after co-culture with Karpas-422. Effector cytokines (Granzyme B, IFN-γ, MIP-1a, perforin, TNF-α, TNF-β); Stimulatory cytokines (GM-CSF, IL-12, IL-15, IL-2, IL-21, IL-5, IL-7, IL-8, IL-9); Chemoattractive cytokines (CCL-11, IP-10, MIP-1b, RANTES); Regulatory cytokines (IL-10, IL-13, IL-22, IL-4, sCD137, sCD40L, TGF-b1); Inflammatory cytokines (IL-17A, IL-17F, IL-1b, IL-6, MCP-1, MCP-4). (F-I) Microwell assay using TROVO, comparing single cell killing rate (F) (n=3 technical replicates; two-tailed Student’s *t*-test), T cell proliferation (G) (n=2 donorsx3 replicates; two-tailed Student’s *t*-test), and tumor killing (H) (n=2 donors x 3 replicates each, two-tailed Student’s *t*-test). (I) Representative images used to calculate data shown in (G) and (H). Each row represents one microwell monitored over 4 days. Each microwell is set up with E:T =20:10. T cell = red. \**P<0.05; **P<0.01; ***P:<0.001; ****P:<0.0001*.

### VHH2-CAR-T eliminated MET+ tumor cells in vitro and in vivo

Since MET overexpression is associated with poor prognosis in many solid tumors, we tested the efficacy of VHH2-CAR-T against different subtypes of solid tumors with known MET overexpression. VHH2-CAR-T demonstrated cytotoxicity against tumor cell lines from TNBC (MDA-MB-231 and LM2), clear cell renal cell carcinoma (786-O), and head and neck squamous cell carcinoma (CAL27 and FaDu) (**Figures 3A and B**).

**Figure 3:**
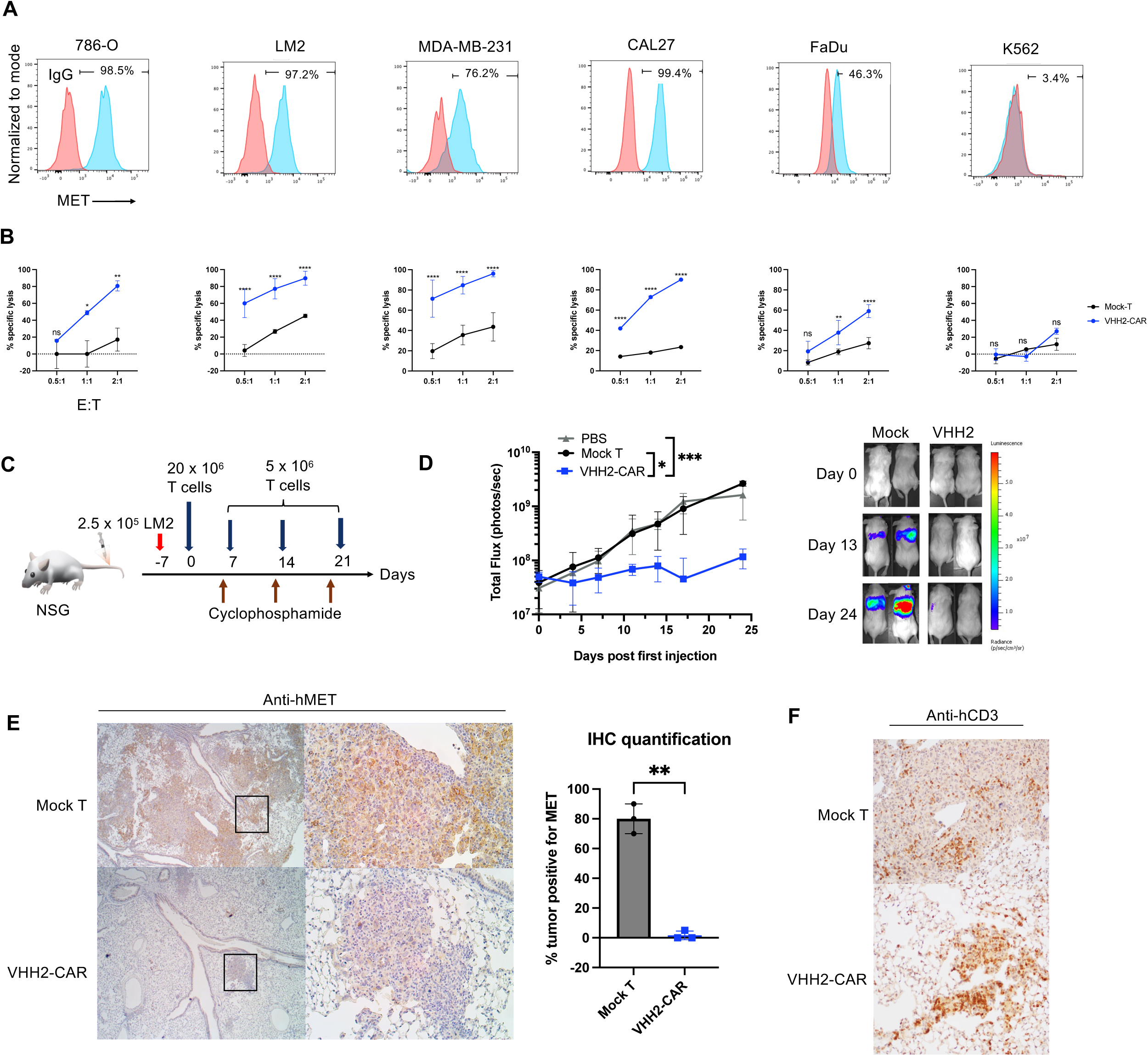
In vitro and in vivo efficacy of VHH2-CAR. (A) MET expression in tumor cell lines. Red = IgG isotype control. (B) Cytotoxicity of VHH2-CAR-T cells at different E:T ratios (n=3 technical replicates; two-way ANOVA). (C) Scheme of in vivo experiment using the LM2 tumor model. (D) BLI of tumor growth in vivo. PBS = 3 animals; Mock T = 2 animals; VHH2-CAR-T = 3 animals. *P* values were calculated using two-way ANOVA. (E) MET IHC, 10x objective on the left and 20x objective on the right. The boxed regions were magnified and shown on the right. The bar graph represents quantifications of MET expression. *P*-value was calculated by two-tailed Student’s *t-*test. (F) CD3 IHC, 20x objective. \**P<0.05; **P<0.01; ***P<0.001; ****P<0.0001*.

Recent clinical trials have shown that repeated CAR-T cell injections exert better tumor control than single CAR-T administration in solid tumors[34,35]; presumably because fresh supplies of T cells help overcome T cell exhaustion in the tumor microenvironment [34]. We tested the in vivo efficacy through weekly injections of fresh mRNA VHH2-CAR-T cells (**Figure 3C**). We focused on the aggressive lung metastatic TNBC model LM2. Luciferase-expressing LM2 tumor cells were injected intravenously (IV) via the tail vein into NSG mice. After 7-10 days, we confirmed tumor growth in bilateral lungs by bioluminescence imaging (BLI). We then injected PBS, 20×10^6^ mRNA VHH2-CAR-T cells or an equal number of mock T cells via the tail vein, followed by three additional weekly injections of 5×10^6^ effector cells or PBS.

Significant reduction in tumor growth was observed in the VHH2-CAR-T group compared to the mock T cell group (**Figure 3D**). Mice were euthanized on day 25 for necropsy. Immunohistochemical (IHC) staining with anti-MET revealed heterogeneous MET expression within the tumor bed in the mock T cell and PBS groups (**Figure 3E**). Although some residual tumor cells were seen in the VHH2-CAR-T group, they were mostly MET negative by IHC (**Figure 3E**); the tumor growth curve also suggested slow growth of these residual tumors (**Figure 3D**). In the VHH2-CAR-T group, IHC for human CD3 showed clear infiltration and aggregation of T cells in the residual tumors, whereas the mock T cell group showed only limited numbers of single scattered T cells (**Figure 3F**). Given the aggressive nature of tumors generally associated with MET overexpression, these findings indicate that VHH2-CAR-T could efficiently control tumor growth by eliminating MET-positive tumor cells.

### VHH2-CAR-T activity is dependent on MET expression level

Given the necropsy findings that MET can be expressed at heterogenous levels in different tumor cells, we next tested the antigen threshold for VHH2-CAR-T activation. First, in a panel of small cell lung cancer cell lines with varying levels of MET expression[23] (**Figure 4A**), we observed that increased MET expression resulted in increased cytotoxicity by VHH2-CAR-T, indicating that targeted tumor lysis was dependent on MET expression level (**Figure 4B-E**).

**Figure 4:**
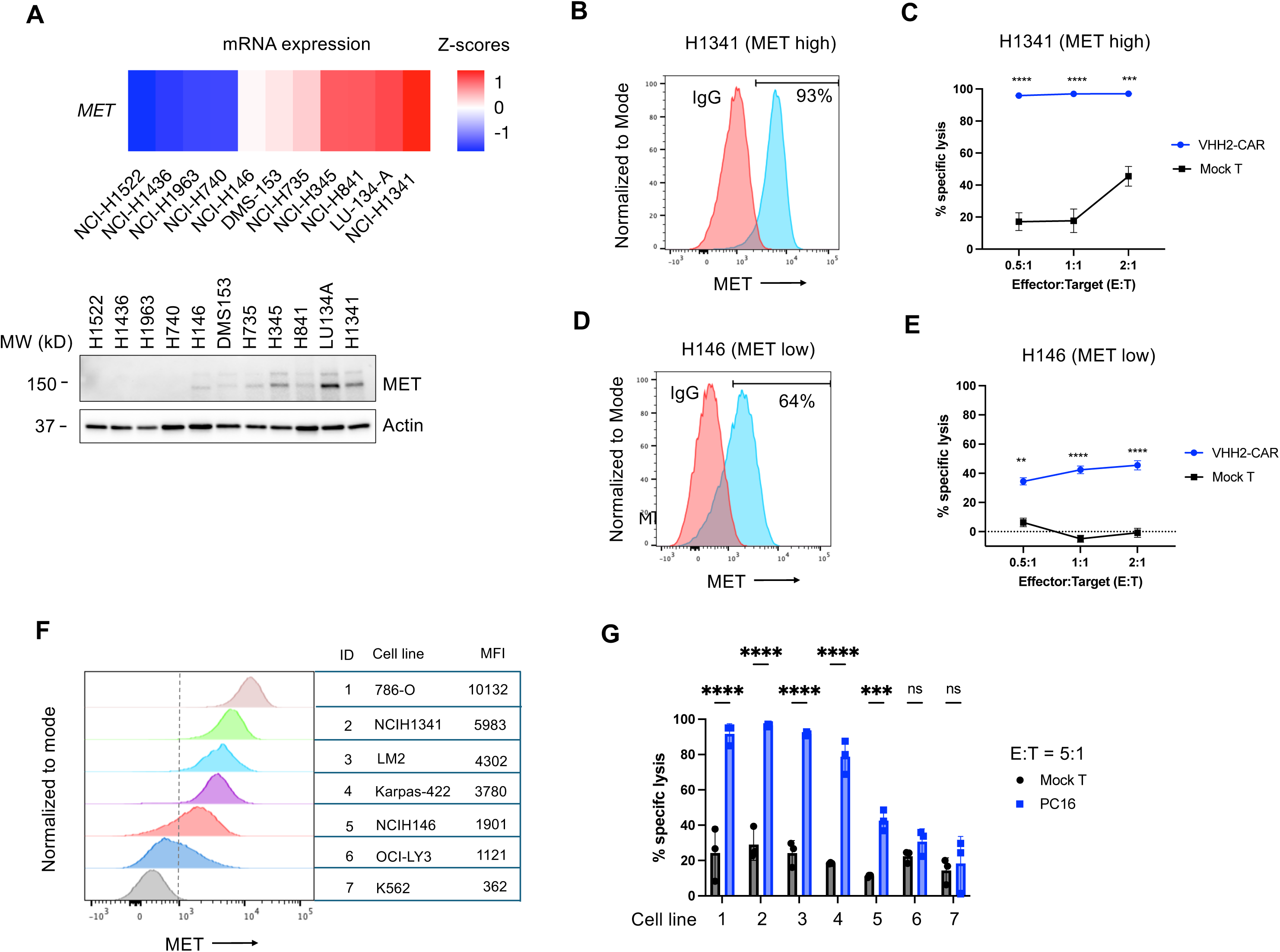
Antigen threshold for VHH2-CAR. (A) mRNA and western blot of MET expression levels in a series of small cell lung cancer cell lines. The mRNA data comes from CellMiner[23]. (B and C) MET expression by flow cytometry and cytotoxicity in NCI-H1341. N= 3 technical replicates in the cytotoxicity assay; *P* values from two-way ANOVA. (D-E) MET expression by flow cytometry and cytotoxicity in NCI-H146. N= 3 technical replicates; *P* values calculated using two-way ANOVA. (F) Histogram of MET expression levels in different cell lines. (G) Cytotoxicity of VHH2-CARs against the same cell lines as in (E). N= 3 technical replicates; *P* values calculated using two-tailed Student’s *t-*test. \**P<0.05; **P<0.01; ***P<0.001; ****P<0.0001*.

To further investigate antigen-dependent cytotoxicity, we compiled and analyzed a series of cell lines with varying levels of MET expression and tested cytotoxicity at a high E:T ratio of 5:1 (**Figure 4F-G**). As expected, co-culture of VHH2-CAR-T with cell lines with high levels of MET expression (786-O, NCIH1341, LM2, and Karpas-422) led to significant tumor cell death ranging from 80-100%. Although the intermediate MET-expressing cell line, NCIH146, still showed significant cell death, the killing by VHH2-CAR-T dropped to approximately 45-50% (against a background of ∼10% mock T cell killing). For the low-MET cell line, OCI-LY3, and MET-negative cell line, K562, VHH2-CAR-T showed cytotoxicity not significantly different from mock T cells.

Taken together, our findings suggest the existence of an antigen threshold level for VHH2-CAR-T activity. In normal tissues, MET is expressed at the highest level in hepatocytes, whose MET expression level is predicted to fall between OCI-LY3 and K562 cell lines, for which we observed limited cytotoxicity and a potential therapeutic window for VHH2-CAR-T cells.

### VHH-CAR-T cells showed low tonic signaling

Since nanobodies generally show high stability[8,12], we tested how incorporation of VHH influences CAR-T cell function compared to MET-CAR-T cells engineered with different sources of antigen binding domains (ABDs).

To facilitate comparison, we adapted a previously reported CAR-Jurkat (CAR-J) assay [24] (**Figure 5A**). We electroporated mRNA encoding CARs with various ABDs into Jurkat cells, co-cultured them with tumor cell lines at various effector to target ratios for 16 hours, and assessed expression of CD69 activation marker by flow cytometry. The diffuse large B-cell lymphoma (DLBCL) cell line, Karpas-422, served as a convenient target for this assay as it expresses high levels of CD19 in addition to MET, thus allowing comparison with CD19-CAR (FMC63) as a positive control (**Supplemental Figure 3**).

**Figure 5:**
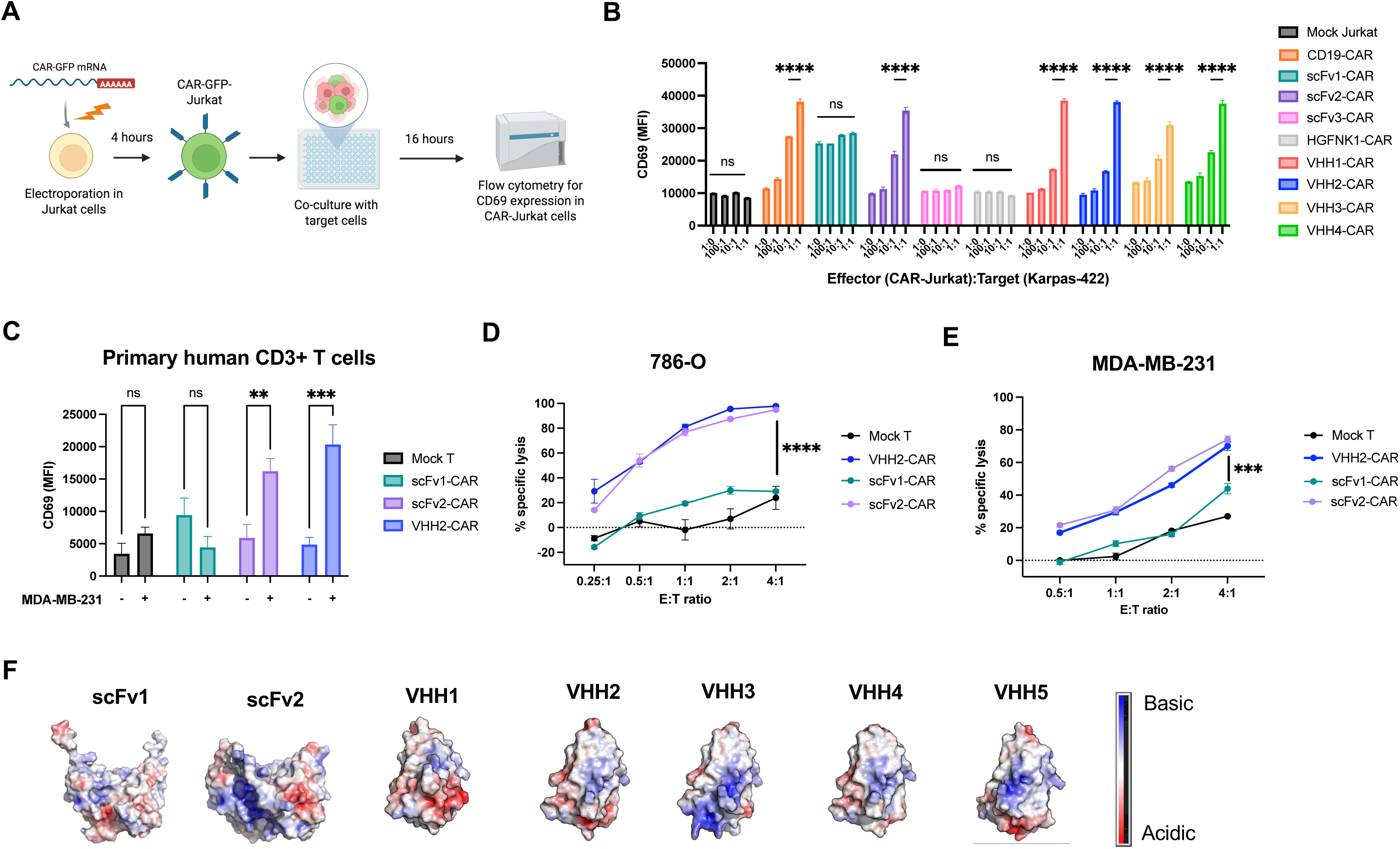
VHH-CARs showed low tonic signaling. (A) Experimental design of CAR-Jurkat assay. (B) Comparison of CD69 expression in CAR-Jurkat cells with varying levels of antigen stimulation. (C) Comparison of CD69 expression in primary human T cells electroporated to express CARs. E:T ratio was 1:1, and co-culture was incubated for 16 hours before staining for flow cytometry. (D-E) Cytotoxicity comparison between VHH2, scFv1, and scFv2 using two different cell lines with different MET expression levels. (F) Electrostatic surface charge of MET binders. Modeling was performed using PyMol. Blue = more basic/positive charge; red = more acidic/negative charge; white = neutral. In experiments shown in B-E, n=3 technical replicates, and *P* values were calculated with two-way ANOVA. \**P<0.05; **P<0.01; ***P<0.001; ****P<0.0001*.

The selected ABD constructs included three different scFvs, one natural ligand, and VHH1-4 (**Figure 5B**). ScFv1 and scFv3-CAR-J were constructed similarly to previously described cell lines [19,20]; scFv2-CAR-J was derived from the monoclonal antibody Onartuzumab (5D5)[36]. Finally, a natural ligand CAR previously shown to mediate cytotoxicity against mesothelioma was derived from the NK1 domain of HGF[21].

As expected, CD19-CAR-J cells showed an antigen-dependent increase in CD69 expression, whereas mock Jurkat cells showed only basal levels of expression at all E:T ratios (**Figure 5B**). The tested scFv-CAR-J cells, however, demonstrated a wide range of antigen-dependent responses. Notably, scFv1-CARJ showed high basal levels of CD69 expression in the absence of Karpas-422, indicating tonic signaling. Indeed, increasing the target cell ratio did not lead to a further increase in CD69 expression, suggesting T cell dysfunction. In contrast, scFv2-CAR-J showed low basal levels and an antigen-dependent increase in CD69. scFv3-CAR-J showed low basal CD69 levels, but for an unknown reason did not show an antigen-dependent response. The ligand-based NK1-CAR-J also failed to show significant CD69 expression upon stimulation. In contrast to the varied behaviors of the tested scFv– and ligand-CAR-J cells, all VHH-CAR-J cells demonstrated low basal levels and an antigen-dependent increase in CD69 expression. The various tonic signaling profiles seen in Jurkat cells could be replicated in co-cultures of primary human T cells and MDA-MB-231 target cells (**Figure 5C**).

To further show that tonic signaling leads to T cell dysfunction, we chose the tonic scFv1 and the low-tonic scFv2 and VHH2 for comparison. As expected, scFv1-CAR-T cells showed only limited tumor killing compared to scFv2– and VHH2-CAR-T cells, both of which showed equivalent tumor killing at various E:T ratios (**Figure 5D and 5E**).

The predisposition of different ABDs for tonic signaling is intriguing. Because previous reports have correlated positive surface charges of ABDs with tonic signaling[37], we modeled the electrostatic surfaces for scFv1, scFv2, and VHH2 (**Figure 5F**). However, structural modeling showed no obvious differences in net surface charges, suggesting that other mechanisms might contribute to tonic signaling (see Discussion).

### Different binders endowed varying functional profiles to CAR

We further tested whether low tonic signaling of VHH affects avidity, cytokine secretion, and tumor killing kinetics. Again, we compared VHH2 with scFv1 and scFv2. All three appear to share similar predicted affinities (5-10 nM). However, structural modeling predicts different binding epitopes on MET; while VHH2 and scFv1 occupy a more membrane-proximal epitope, scFv2 binds to a more membrane-distal one (**Figure 6A**).

**Figure 6:**
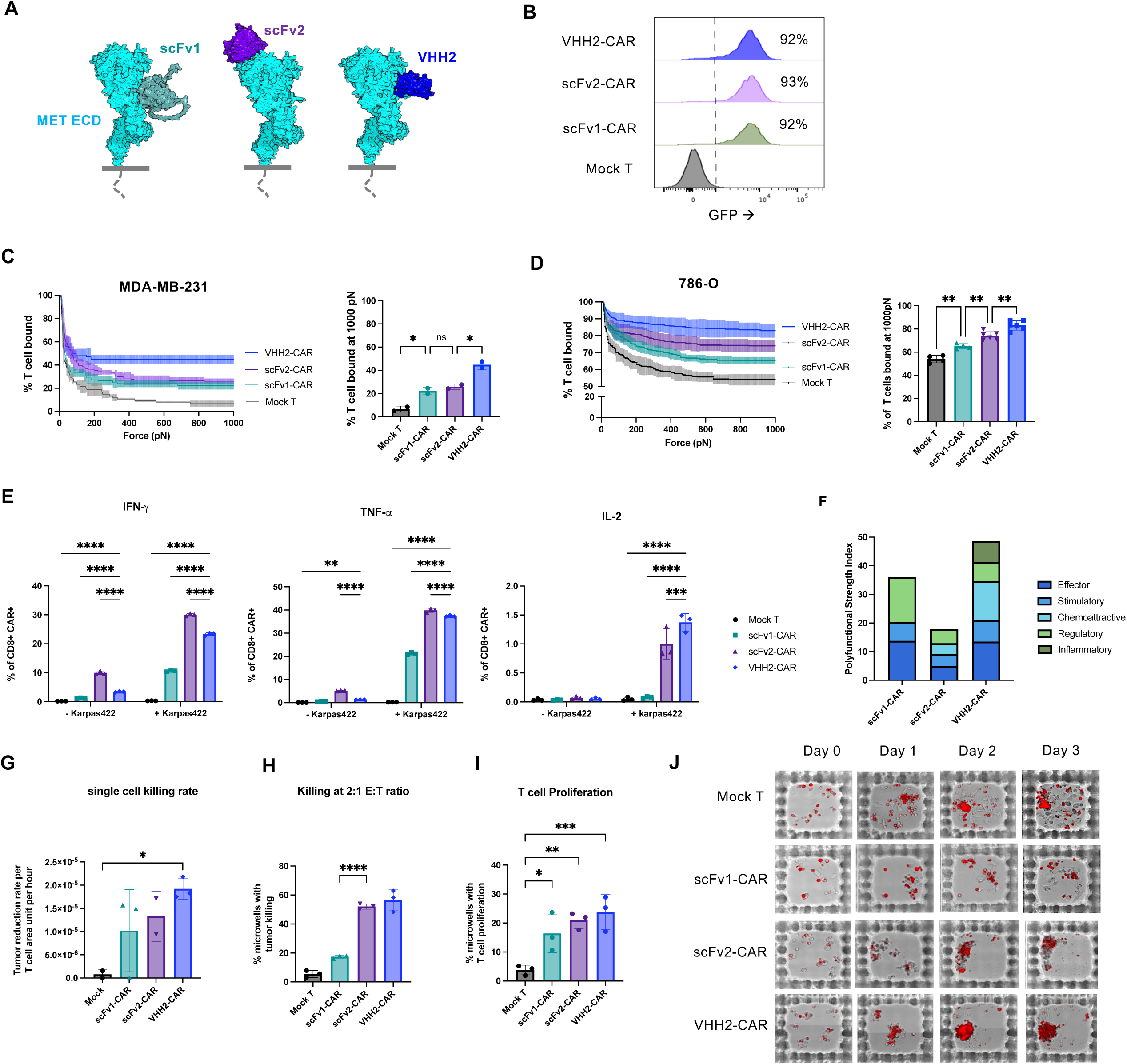
Avidity and kinetics comparisons of CARs. (A) Different binding epitopes predicted by AlphaFold3 for VHH2, scFv1, and scFv2. (B) Expression of compared CARs. (C) Avidity measurement by Lumicks z-Movi using MDA-MB-231 as target cells. N = 3 technical replicates, with *P* values from two-tailed Student’s *t-*test. (D) Avidity measurement against the 786-O cell line. N = 4-5 technical replicates; *P* values from two-tailed Student’s *t-*test. (E) Intracellular cytokine production by flow cytometry. Cells were stained before and 16-hours after stimulation by Karpas-422. N=3 technical replicates; *P* values from two-way ANOVA. (F) Polyfunctional strength index by IsoPlexis. Effector cytokines (Granzyme B, IFN-γ, MIP-1a, perforin, TNF-α, TNF-β); Stimulatory cytokines (GM-CSF, IL-12, IL-15, IL-2, IL-21, IL-5, IL-7, IL-8, IL-9); Chemoattractive cytokines (CCL-11, IP-10, MIP-1b, RANTES); Regulatory cytokines (IL-10, IL-13, IL-22, IL-4, sCD137, sCD40L, TGF-b1); Inflammatory cytokines (IL-17A, IL-17F, IL-1b, IL-6, MCP-1, MCP-4). (G-I) Hydrogel-based microwell co-culture assay by TROVO, showing single cell killing rate (G, 1:10 E:T), killing (H, 20:10 E:T), and T cell proliferation (I, 20:10 E:T). N=3 technical replicates; *P* values using two-tailed Student’s *t-*test. (J) Representative images of microwell co-culture at 20:10 E:T. Each row represents one microwell monitored over time. \**P<0.05; **P<0.01; ***P<0.001; ****P<0.0001*.

We first compared how the three ABDs influence avidity, expecting roughly equal avidity given their equivalent affinities. Interestingly, despite similar expression levels of all constructs (**Figure 6B**), VHH2 consistently showed stronger avidity than scFv2, which was greater than scFv1 for two different tumor cell lines (**Figures 6C-D**). The low avidity of scFv1 could be related to factors that also contributed to its tonic signaling. The stronger avidity of VHH2 over scFv2 was unclear, but one possible explanation could be related to VHH2’s binding to a more membrane-proximal epitope, as targeting membrane-proximal epitopes has been shown to stabilize the immune synapse[38,39].

In terms of cytokine production, both scFv2 and VHH2 showed higher levels of IFN-γ, TNF-α, and IL-2 after target stimulation than scFv1 (**Figure 6E** and **Supplemental Figure 4**). Among the three constructs, scFv2 was noted to have the highest baseline cytokine secretion in the absence of antigens (especially IFN-γ), suggesting some tonic signaling in scFv2. Using IsoPlexis, VHH2 demonstrated the highest polyfunctional strength index (PSI) in vitro (**Figure 6F**), including the highest effector, stimulatory, and chemoattractive and lower regulatory cytokine levels. Although there were increased inflammatory cytokines detected by IsoPlexis for VHH2, by ELISA VHH2 produced lower levels of proinflammatory cytokines such as IL-6, IL-8, IL-10, IL-1β, and granzyme A (**Supplemental Figure 4**). The reason for this difference is unclear but might be due to different assay conditions.

We also examined the kinetics of tumor cell killing using a microwell platform monitored over four days (as in Figure 2F-I). Although not statistically significant, VHH2-CAR-T cells trended to have the highest single-cell killing rate (**Figure 6G**), the highest percentage of microwells with tumor killing (**Figure 6H**), as well as the highest T cell proliferation (**Figure 6I**). Taken together, these data suggest diverse behaviors of different MET-CAR-T cells, and that VHH2-CAR-T cells have high avidity, potent immune cytokine production including polyfunctionality, and rapid tumor killing kinetics.

### VHH2-CAR-T showed potent in vivo efficacy in comparison with scFv2-CAR-T

Emerging evidence suggests that avidity correlates with in vivo and clinical efficacy more so than in vitro cytotoxicity[31]. We tested this hypothesis by comparing VHH2 and scFv2 in vivo given their overall similar in vitro cytotoxicity but differing avidities. Since multiple high doses of VHH2-CAR-T cells appeared to provide potent tumor control (**Figure 3**), we performed a CAR-T cell stress test by reducing the weekly dose to 2×10^6^ CAR-T cells per IV injection and monitored tumor growth over a longer time period (**Figure 7A**).

**Figure 7:**
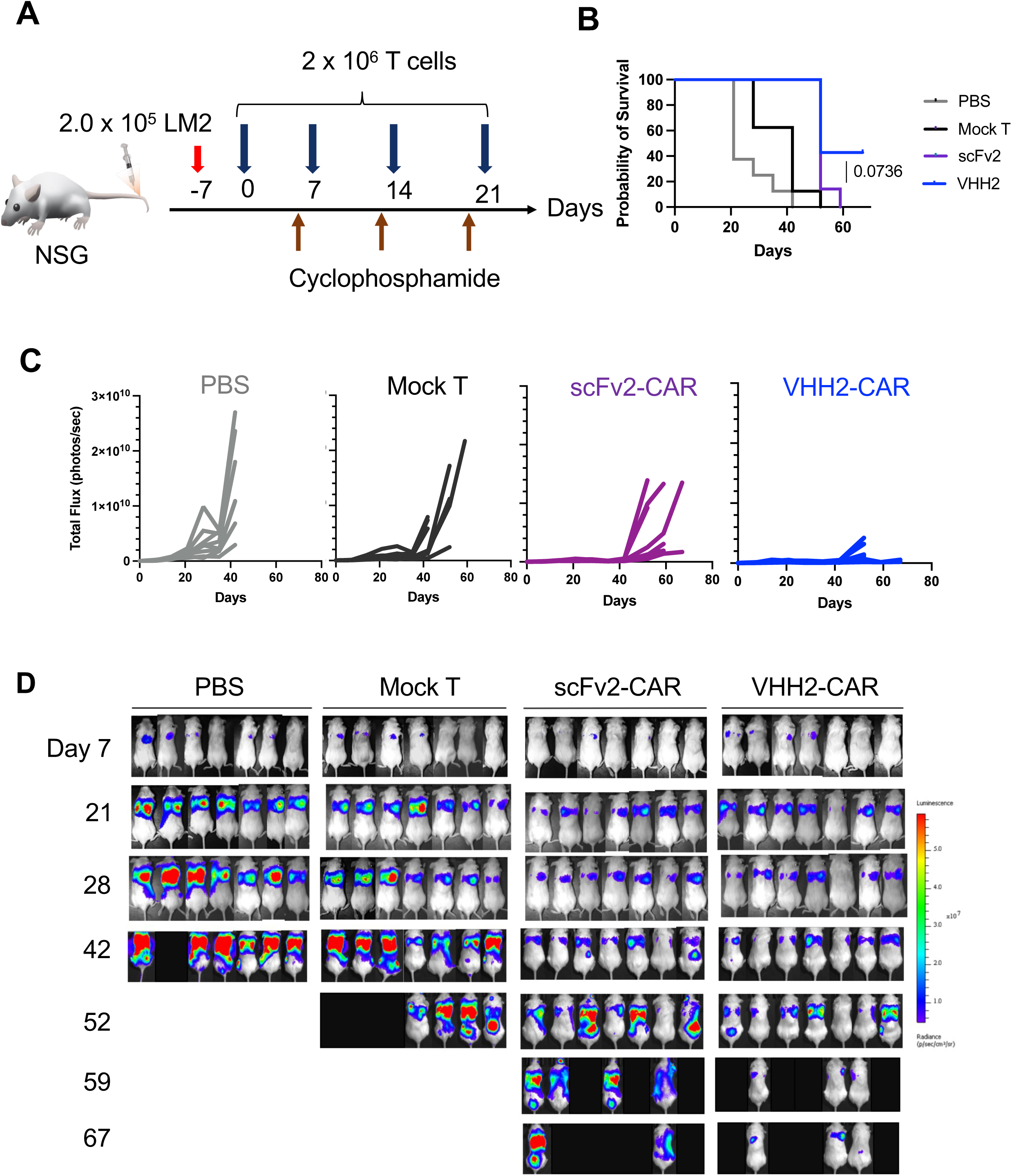
In vivo comparison between scFv2 and VHH2-CAR. (A) Experimental design of in vivo efficacy. (B) Kaplan-Meier survival curve using a predefined tumor burden threshold of 1×10^9^ photons/sec as the survival endpoint for analysis purposes. *P*-value shown for VHH2-CAR versus scFv2-CAR comparison. (C) Tumor growth curve by BLI imaging. Each group = 7 mice; each line = 1 mouse. (D) Representative images of BLI.

Both VHH2-CAR-T and scFv2-CAR-T groups showed significantly better survival than mock T cells and PBS, and VHH2-CAR-T conferred slightly longer survival than scFv2-CAR-T (close to statistical significance) (**Figures 7B-D**). There was a significant reduction in tumor growth rate in the VHH2 and scFv2 groups compared to saline or mock T cells (**Figures 7C-D**). Interestingly, after the last CAR-T cell injection at day 21, many animals in the VHH2 group showed lower tumor burdens than the scFv2 group. Three animals in the VHH2 group showed continuous tumor control until approximately day 67. Overall, there were also fewer extrapulmonary metastases in the surviving mice in the VHH2 group compared to the scFv2 group (**Figure 7D**). Collectively, our in vivo model indicates that VHH2-CAR-T cells are as effective as scFv2-CAR-T cells with modestly enhanced tumor control, likely due to more complete elimination of MET-positive tumor cells by the stronger avidity VHH2.

Although there was no overall survival difference between VHH2– and scFv2-CAR-T cells, some of the mice appeared to die due to factors other than tumor burden thus complicating survival curve analysis. On-target/off-tumor cytotoxicity (OTOT) was unlikely as the VHH2 does not bind murine MET[40], and histology showed no obvious evidence of tissue damage. Other etiologies including cytokine storm or development of early GVHD were also possible.

## DISCUSSION

We used an array of functional assays to comprehensively characterize nanobody-based CAR-T against MET-positive solid tumors. Among a panel of VHH, we found that VHH1 and VHH2 with intermediate avidity mediated the best killing especially at low E:T ratios. The CD28 costimulatory domain enhanced VHH2-CAR-T cell in vitro cytotoxicity and effectively controlled tumor growth in vivo. Despite their potency, VHH2-CAR-T cells displayed an antigen threshold that could potentially discriminate between tumor and normal tissues. Compared to the more variable behaviors of scFv-CAR-T cells, VHH-CAR-T cells showed low tonic signaling, high avidity, a favorable cytokine profile, and rapid tumor killing kinetics – features that likely contributed to prolonged tumor control in vivo despite the transient nature of the injected mRNA CAR-T cells.

This study highlights construct design as a crucial step in optimizing CARs. We examined the fundamental aspects of changing the ABD with VHH as a comparison to scFv or ligand-based approaches. Nanobodies are known for their small size and high stability [8–10]. We demonstrated that all our tested VHHs have low tonic signaling, whereas the selected scFv were more variable, with scFv1 being the most prominent example of a tonic signaling CAR. Interestingly, structural modeling of scFv1 did not indicate excessive surface positive charges that were previously reported to increase tonic signaling potential[37]. However, other mechanisms, such as hydrophobic interactions and light-chain-heavy-chain mispairing are also known to occur in scFv and might contribute to the tonic signaling of scFv1[5]. Although not without limitations (such as the need for humanization), VHHs might generally offer an alternative solution for some tonic signaling CARs [4].

We found that the two intermediate avidity VHH1 and VHH2 were more efficacious at eliminating tumors than either the higher avidity VHH4/VHH5 or the lower avidity VHH3. There is a complicated relationship between avidity and cytotoxicity—some prior publications suggest that increased avidity leads to better CAR-T cell performance[41,42], while others indicate that decreased avidity is preferable[38,43]. One potential explanation is that CAR-T cells with high avidity have low *Koff* and therefore prolonged engagement with tumor cells, which can lead to activation-induced cell death and compromised tumor killing[38,43]; on the other hand, excessively low avidity translates to an unstable immune synapse. In another context, VHH2-28z showed a trend toward higher avidity and cytotoxic function than VHH2-BBz, likely due to the predisposition of CD28 to dimerize and thus promote multivalent interactions[32,44,45]. Finally, the higher avidity of VHH2 compared to scFv2 could potentially be related to VHH2’s binding to a more membrane-proximal epitope and correlated with slightly better tumor killing kinetics and tumor control in vivo. These findings reflect the multifactorial nature of avidity, which depends on CAR-target antigen affinity, binding valency, and CAR-antigen orientation, among other factors. Future studies will benefit from a better understanding of the factors that influence the avidity-CAR function relationship.

Although the CD28 costimulatory domain led to higher and faster killing as shown here, other studies have demonstrated earlier T cell exhaustion than 4-1BB[32]. However, we did not observe notable differences in PD-1 and LAG3 expression between VHH2-28z versus VHH2-BBz, perhaps due to short duration of mRNA CAR expression. For targeting solid tumors, it might be advantageous to couple CARs with the CD28 costimulatory domain in an mRNA format to achieve immediate and maximal tumor killing without the development of T cell exhaustion, as outlined in a recent review[34].

VHH2-CAR-T cells appeared to have an antigen threshold for activation and thus a possible therapeutic window. Encouragingly, a phase 1 trial of a MET CAR using the same scFv2 (5D5) demonstrated safety with mostly clinically manageable grade 1-2 cytokine release syndrome (CRS)[46]. Given that CAR-T cells generally require a high number of target antigens to be present to become activated, therapeutic windows are likely present between normal tissues and MET-overexpressed tumors, as shown here and illustrated elsewhere[47]. One future direction includes exploring logic-gated CAR design to further minimize on-target/off-tumor toxicity.

Despite the short duration of our mRNA CARs, some VHH2-treated mice displayed remarkably prolonged tumor control until day 67, even after cessation of CAR-T cell injections on day 21. This prolonged tumor control was likely due to the ability of the high avidity and polyfunctional VHH2-CAR-T cells to kill most of the MET-positive cells, which were presumably more proliferative than the MET-negative cells within the same tumor. One can imagine a future strategy against solid tumors that would deploy potent effector like VHH2-CAR-T cells to target a surface antigen related to tumor aggressiveness (e.g. MET) to achieve prolonged tumor control before bridging to other therapies.

The use of mRNA in the current study offered many advantages but also some limitations. The main advantages included uniform CAR expression and high T cell viability, which facilitated construct comparisons in multiple assays. Looking forward, mRNA represents a particularly promising platform for CAR-T cell reprogramming given the current progress in developing in vivo nanoparticle delivery systems[48]. However, in some experiments, such as the microwell co-culture and xenograft mouse models, it would have been beneficial to use a persistent CAR system so that one could monitor long-term tumor control and T cell functions. In summary, we have shown the efficacy of a novel class of VHH-CAR-T cells against MET with promising in vitro attributes as well as strong tumor control in vivo. The VHH-CAR-T cell system warrants further preclinical study and development, including VHH humanization and further construct optimization.

### Declarations

#### Data availability statement

All data relevant to the study are included in the article or uploaded as supplementary information.

#### Ethics statements

Patient consent for publication: Not applicable.

#### Ethics approval

This study did not include human subjects. All animal studies were approved by the Yale Institutional Animal Care and Use Committee.

#### Author contributions

P-H.C. and S.G.K. designed all the experiments and wrote the manuscript. P-H.C. designed and performed all the experiments. Q.L., S.D., D.K.S., and P-C.C. contributed to data figures and experiments. R.R. and N.C. contributed to molecular biology and cloning of different CAR constructs. M.N. and A.A. contributed cell lines and performed western blot and mRNA analysis. Y.L. and Q.Y. helped design and perform the *in vivo* experiment. All authors reviewed the manuscript.

## Acknowledgments

The research was performed as part of the Yale Pathology Physician-Scientist training track. We also appreciate the support from the K12 Yale Immuno-Oncology Training Program at Yale Cancer Center. We thank the Yale Animal Research Facility, for excellent technical support. We also thank the Yale Flow Cytometry Core Facility for maintaining and making available the cytometers used in the research. The Core is supported in part by an NCI Cancer Center Support Grant # NIH P30 CA016359. We also thank Keck Microarray Shared Resource (KMSR) at Yale University for providing the necessary services using the IsoSpark system, which is funded in part by the National Institutes of Health instrument grant 1S10OD034429-01.

## SUPPLEMENTAL FIGURES

**Supplemental figure 1:** mRNA CAR manufacturing through electroporation. (A) mRNA CAR design. (B) RNA gel showing examples of successful mRNA synthesis and poly(A) tail incorporation. (C) CAR expression was dose-dependent as shown by GFP. (D) Time course of GFP decay after mRNA electroporation in T cells.

**Supplemental figure 2:** Intracellular cytokine assessment. (A) Gating strategy. (B) Example flow cytometry plot of cytokine production.

**Supplemental figure 3:** Lack of significant exhaustion marker expression in VHH2-28z. (A) Representative flow cytometry plot of PD-1 versus TIM3 and PD-1 versus LAG3 expression. CAR-T cells were co-cultured with Karpas-422 for 48 hours before harvesting for flow cytometry. (B) Comparison of PD-1 and LAG3 expression, and PD-1/TIM3 coexpression between mock T, VHH2-28z, and VHH2-BBz T cells. N=3 technical replicates; *P* values from two-way ANOVA. \**P<0.05; **P<0.01; ***P<0.001; ****P:<0.0001*.

**Supplemental Figure 4:** Karpas-422 cell line with MET and CD19 expression. (A) Flow cytometry demonstrating expression of both antigens. Red curve = IgG isotype control. (B) Confirmation of cytotoxicity of different MET CAR-T cells and CD19-CAR-T cells against Karpas-422. N = 3 technical replicates, with *P* values calculated from two-tailed Student’s *t-*test. \**P<0.05; **P<0.01; ***P<0.001; ****P:<0.0001*.

**Supplemental Figure 5:** Bulk cytokine secretion by ELISA between various CAR constructs against MDA-MB-231. Panel A shows selected effector cytokines, while panel B shows selected cytokines known to mediate CAR T cell related toxicity and inflammatory responses.

