## Supplemental Figures for "Nanobody MET CAR-T cells show efficacy in solid tumors"

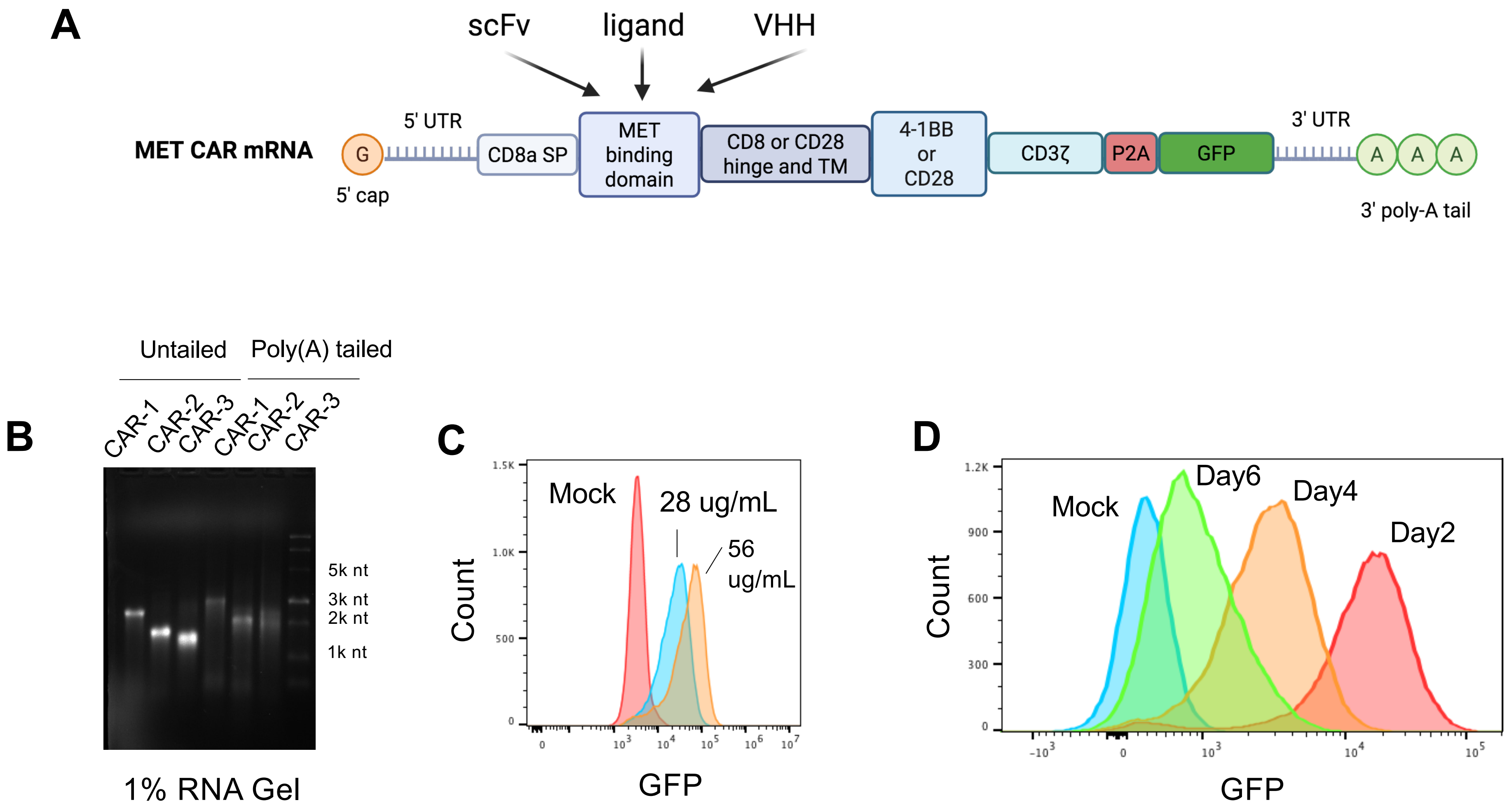


Chen, et al., Supplemental Figure 1

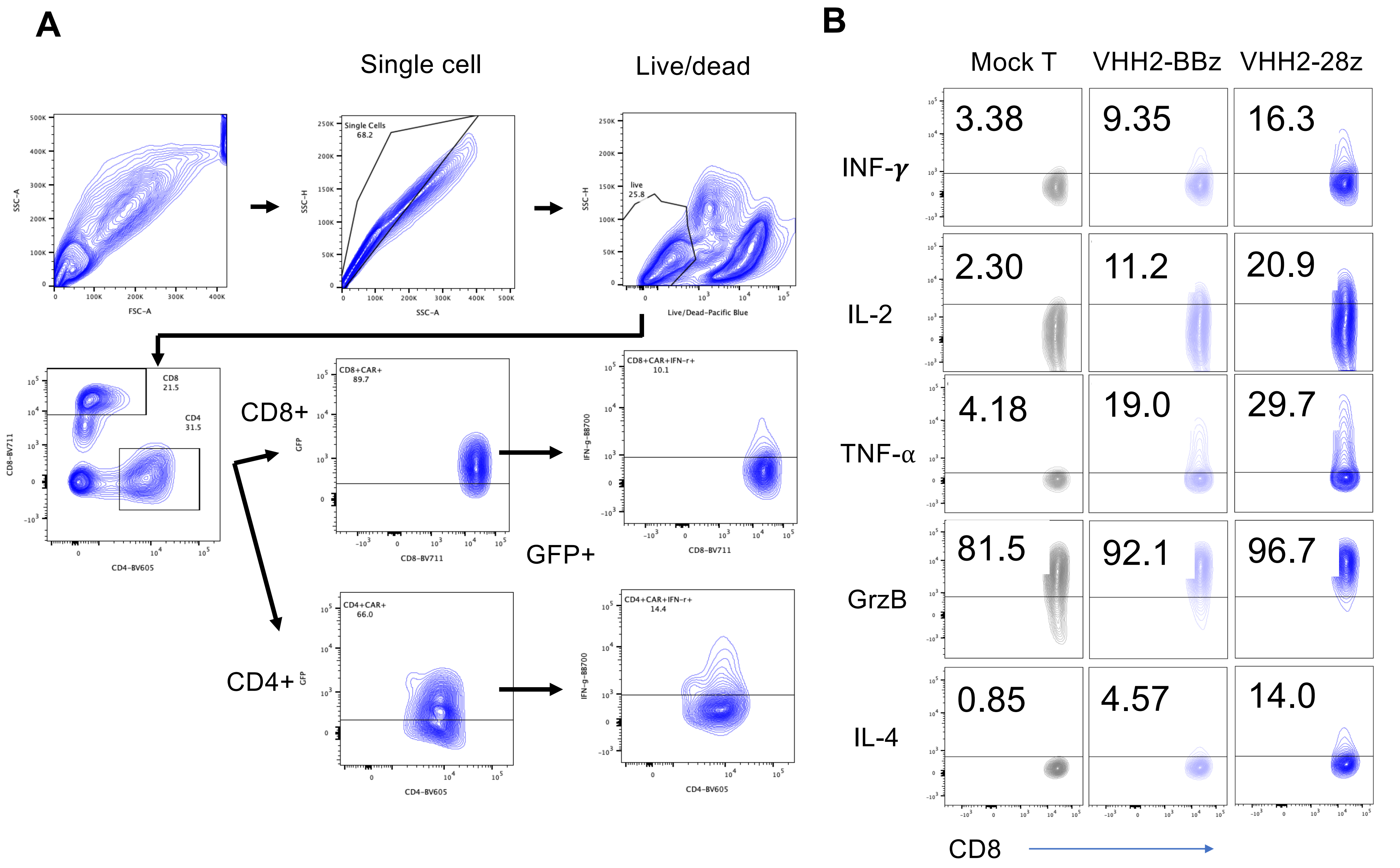


Chen, et al., Supplemental Figure 2

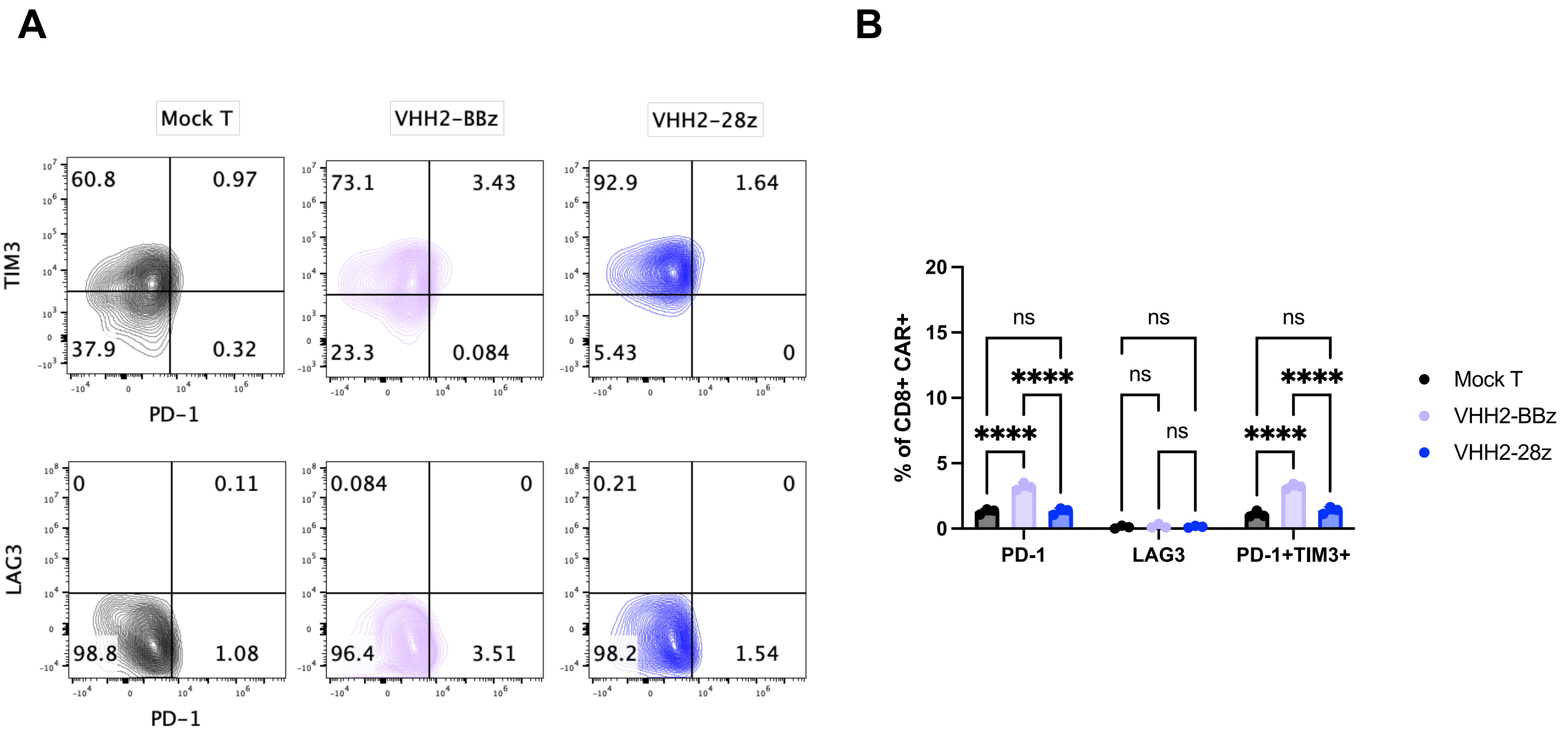


Chen, et al., Supplemental Figure 3

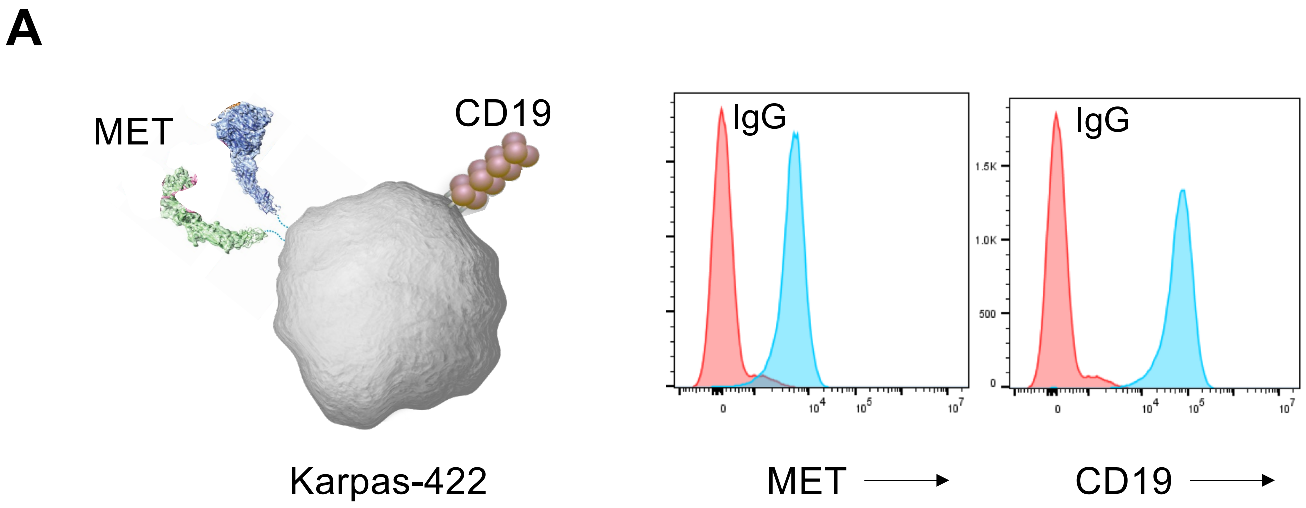


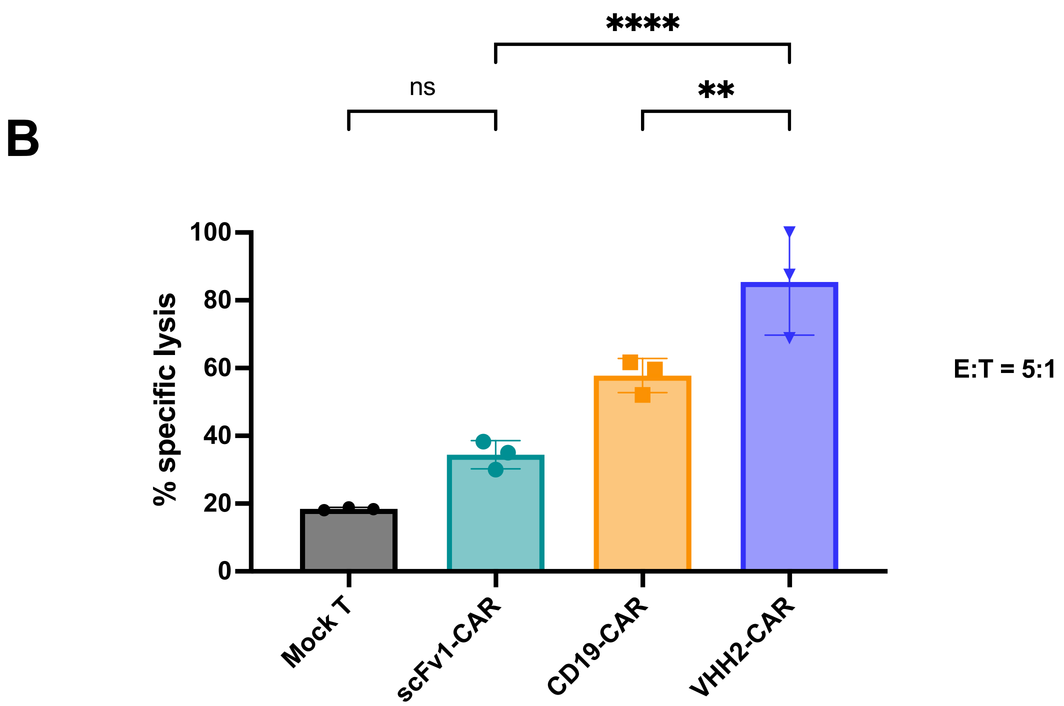


Chen, et al., Supplemental Figure 4

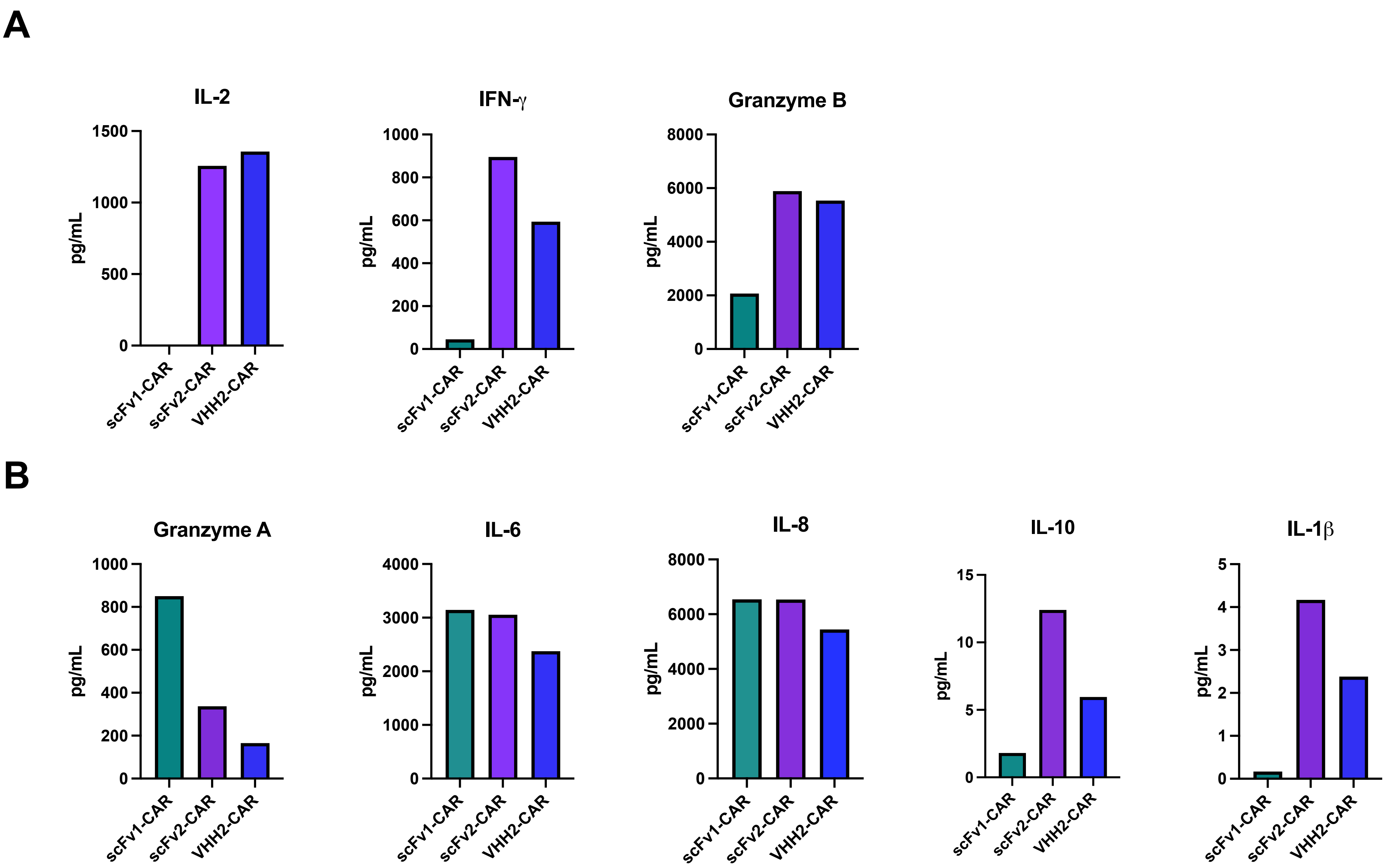


Chen, et al., Supplemental Figure 5
